# The impact of partitioning on phylogenomic accuracy

**DOI:** 10.1101/023978

**Authors:** Diego Darriba, David Posada

## Abstract

Several strategies have been proposed to assign substitution models in phylogenomic datasets, or partitioning. The accuracy of these methods, and most importantly, their impact on phylogenetic estimation has not been thoroughly assessed using computer simulations. We simulated multiple partitioning scenarios to benchmark two *a priori* partitioning schemes (one model for the whole alignment, one model for each data block), and two statistical approaches (hierarchical clustering and greedy) implemented in PartitionFinder and in our new program, PartitionTest. Most methods were able to identify optimal partitioning schemes closely related to the true one. Greedy algorithms identified the true partitioning scheme more frequently than the clustering algorithms, but selected slightly less accurate partitioning schemes and tended to underestimate the number of partitions. PartitionTest was several times faster than PartitionFinder, with equal or better accuracy. Importantly, maximum likelihood phylogenetic inference was very robust to the partitioning scheme. Best-fit partitioning schemes resulted in optimal phylogenetic performance, without appreciable differences compared to the use of the true partitioning scheme. However, accurate trees were also obtained by a “simple” strategy consisting of assigning independent GTR+G models to each data block. On the contrary, leaving the data unpartitioned always diminished the quality of the trees inferred, to a greater or lesser extent depending on the simulated scenario. The analysis of empirical data confirmed these trends, although suggesting a stronger influence of the partitioning scheme. Overall, our results suggests that statistical partitioning, but also the *a priori* assignment of independent GTR+G models, maximize phylogenomic performance.

## Introduction

Statistical inference of phylogenetic trees from sequence alignments requires the use of probabilistic models of molecular evolution (Felsenstein, 2004). It is well established that the choice of a particular model of molecular evolution can change the results of the phylogenetic analysis, and not surprisingly, one of the most active areas of research in phylogenetics in recent years has been the development of more realistic models of nucleotide, codon and amino acid substitution/replacement, together with the implementation of statistical methods for the selection of best-fit models for the data at hand (Joyce and Sullivan, 2005; Posada, 2012).

A key aspect in the development of these models has been the consideration of the heterogeneity of the substitution process among sites. Several mixture models have been proposed that assign each site within a locus a probability of evolving under a given rate (Yang, 1994), substitution pattern (Lartillot and Philippe, 2004; Pagel and Meade, 2004), or both (Wu *et al.*, 2013). In particular, a discrete gamma distribution to consider rate variation among sites (Yang, 1996) is used nowadays in practically any phylogenetic analysis. A different approach to account for the heterogeneity of the substitution process consists of defining *a priori* groups of sites (so called partitions) that evolve under the same substitution model, like for example the set of 1st, 2nd or 3rd codon positions in protein-coding sequence alignments (Shapiro *et al.*, 2006) or distinct protein domains (Zoller *et al.*, 2015).

At the genomic scale the heterogeneity of the substitution process becomes even more apparent that at the single-gene scale, as different genes or genomic regions can have very different functions and evolve under very different constraints (Arbiza *et al.*, 2011). Multilocus substitution models that consider distinct models for different partitions of the data assumed to evolve in an homogeneous fashion have been proposed under the likelihood (Ren *et al.*, 2009; Yang, 1996) and Bayesian (Nylander *et al.*, 2004; Suchard *et al.*, 2003) frameworks. In this case, different loci (or loci by codon position) are typically considered as distinct partitions by default, without further justification. However, a number of empirical studies have demonstrated that different partitioning schemes can affect multilocus phylogenetic inference, including tree topology, branch lengths and nodal support (Brandley *et al.*, 2005; Kainer and Lanfear, 2015; Leavitt *et al.*, 2013; Powell *et al.*, 2013; Ward *et al.*, 2010), with maximal differences occurring when whole datasets are treated as single partitions (i.e., unpartitioned). Using computer simulations, Brown and Lemmon (2007) showed that both over and particularly under-partitioning can lead to inaccurate phylogenetic estimates.

If the partitioning scheme can affect phylogenetic analysis, we should try to identify the best-fit partitioning scheme for the data at hand. In principle, predefined partitioning schemes might not be included within the optimal ones, and some statistical model selection procedure needs to be implemented to justify the choice of a particular partitioning scheme, just as it happens when finding the best-fit model of evolution for a single locus (Posada and Crandall, 2001). Unfortunately, the number of partitioning schemes for a multilocus data set can be huge, ranging from considering that a single model fits the whole alignment to assigning a different model to each site/region/gene, and until very recently in practice model selection in phylogenomics was restricted to the comparison of a fixed number of alternative partitions in relatively modest data sets, often using Bayes factors (Bao *et al.*, 2007; Brandley *et al.*, 2005; Brown and Lemmon, 2007; Castoe and Parkinson, 2006; Fan *et al.*, 2011; McGuire *et al.*, 2007; Nylander *et al.*, 2004; Pupko *et al.*, 2002);but see Li *et al.* (2008). Opportunely, the release of PartitionFinder (Lanfear *et al.*, 2012, 2014) made a big difference in this regard, facilitating the automatic statistical selection of partitioning schemes for relatively large multilocus data sets. For this task, PartitionFinder uses combinatorial optimization heuristics like clustering and greedy algorithms, building up on previous ideas raised by Li *et al.* (2008). Also, Wu *et al.* (2013) recently described a sophisticated Bayesian approach for the identification of optimal partitioning scheme, but its heavy computational requirements seem to have prevented its general use. While automated statistical model choice procedures have been shown to result in partitioning schemes with a better fit in real data, often resulting in distinct tree topologies when compared to unpartitioned schemes (Kainer and Lanfear, 2015; Wu *et al.*, 2013), the accuracy of these inferences has not been thoroughly assessed. In order to fill this gap we present here a computer simulation study designed to evaluate (i) the precision of the best-fit multilocus partitioning schemes identified by PartitionFinder and by a new tool for multilocus model selection developed by us, called PartitionTest, and (ii) the accuracy of the phylogenetic trees derived from best-fit and *a priori* partitioning schemes. In this article we evaluate the accuracy of PartitionTest and PartitionFinder under different conditions representing biologically realistic scenarios, including rate variation among loci and lineages, non-homogenous data blocks, and large data sets. In addition, we also analyze some of the real datasets previously used in the evaluation of PartitionFinder. Our results suggest that best-fit partitioning schemes can lead to accurate trees, but also that the *a priori* assignment of independent GTR+G models to each locus performs equally well.

## Results

### Simulation 1: multi-gene phylogenetics

The greedy strategy implemented in PartitionTest (PT-G) recovered most often the true partitioning scheme (*PPR* = 0.305), followed by the greedy strategy implemented in PartitionFinder (PF-G) (*PPR* = 0.255) and the hierarchical clustering implemented in PartitionTest (PT-C) (*PPR* = 0.200) (table 1). The hierarchical clustering implemented in PartitionFinder (PF-C) did much worse (*PPR* = 0.013). PT-C, PT-G, PF-G recovered accurate partitioning schemes, with RI (Rand, 1971) values above 0.93. PF-C performed clearly worse (*RI* = 0.852), also evident from its low ARI (Hubert and Arabie, 1985) values. In general, the hierarchical clustering algorithms overestimated the number of partitions while the greedy algorithms underestimated it. The hierarchical clustering algorithms were several times faster than the greedy algorithms. Overall, PartitionTest was on average 2.6 and 1.5 times faster finding the optimal partition than PartitionFinder, for the greedy and hierarchical clustering algorithms, respectively.

**Table 1.** Partitioning and phylogenetic accuracy for Simulation 1 (multi-gene phylogenetics).

|  |  | K=1 | K=T | K=N | PT-C | PT-G | PF-C | PF-G |
| --- | --- | --- | --- | --- | --- | --- | --- | --- |
| Partitioning accuracy | PPR | N/A | N/A | N/A | 0.2 | 0.305 | 0.013 | 0.255 |
|  | RI | N/A | N/A | N/A | 0.97 | 0.931 | 0.852 | 0.951 |
|  | ARI | N/A | N/A | N/A | 0.775 | 0.696 | 0.031 | 0.771 |
|  | Kdiff | N/A | N/A | N/A | 2.011 | -1.71 | 13.682 | -1.773 |
|  | Kmse | N/A | N/A | N/A | 29.294 | 5.855 | 297.321 | 5.761 |
| Phylogenetic accuracy | PTR | 0.787 | 0.89 | 0.89 | 0.892 | 0.842 | 0.885 | 0.82 |
|  | RF | 0.018 | 0.007 | 0.007 | 0.007 | 0.012 | 0.007 | 0.013 |
|  | BS | 0.019 | 0.006 | 0.01 | 0.007 | 0.006 | 0.011 | 0.006 |
| Average Run Time |  | N/A | N/A | N/A | 01:20:25 | 05:25:50 | 01:59:00 | 14:31:20 |

NOTE.— Different partitioning strategies were evaluated: a single partition (K=1), the “true” partitioning scheme (K=T), each data block as a GTR+G partition (K=N), PartitionTest using the Hierarchical Clustering (PT-C) and Greedy (PT-G) algorithms, and PartitionFinder using the Hierarchical Clustering (PF-C) and Greedy (PF-G) algorithms (all of them assuming independent branch lengths). The Greedy algorithms were used only for simulation replicates with up to 20 partitions (1,000 replicates). The accuracy of the selected partitions was evaluated by the number of times the exact true partitioning scheme was identified (PPR = Perfect Partitioning Recovery), the Rand index (RI) and the adjusted Rand index (ARI). The accuracy of the RAxML trees inferred from the selected partitions was evaluated with the average Robinson-Foulds distance (RF) (scaled per branch), a measurement of the number of times the exact true topology was estimated (PTR = Perfect Topology Recovery), and the average Branch Score difference (BS) (scaled per branch). The average time required to identify the optimal partitioning scheme (Average Run time) was measured in hours, minutes and seconds.

Most strategies performed also well recovering the exact true topology (*P T R*, average perfect topology recovery = 0.820−0.890), in particular when using FT-C. The largest differences were observed when a single partition was assumed to underlie the data (K=1), which resulted in an *P T R* of 0.787. The average RF distances to the true topologies were very small (*RF* = 0.007−0.013) except when K=1, which performed worse (*RF* = 0.018) (table 1). The average number of distinct topologies per replicate across methods was 1.31. Regarding the branch lengths, PT-C, PT-G, PF-G performed as well as using the true partitioning scheme (K=T), while PF-G, K=N, and especially K=1, did worse.

### Simulation 2: mosaic data blocks

In this case, where sites inside the simulated data blocks evolved under two different models, there is not a true partitioning scheme so only the accuracy of the trees inferred from the selected partitioning scheme was evaluated. The different strategies did well recovering the exact true topology (*P T R ≥* 0.827), although K=1 did slightly worse (*P T R* = 0.787) (table 2). The average RF distances were larger than in the previous simulation but still reasonably small (*RF* = 0.012−0.014), with K=1 doing slightly worse again (RF=0.018). The average number of distinct topologies per replicate across methods was 1.02. Branch lengths estimate were quite accurate (*BS* = 0.014−0.020), with the greedy algorithms performing best. In this simulation PartitionTest was on average 2.1 and 2.0 times faster finding the optimal partition than PartitionFinder, for the greedy and hierarchical clustering algorithms, respectively.

**Table 2.**
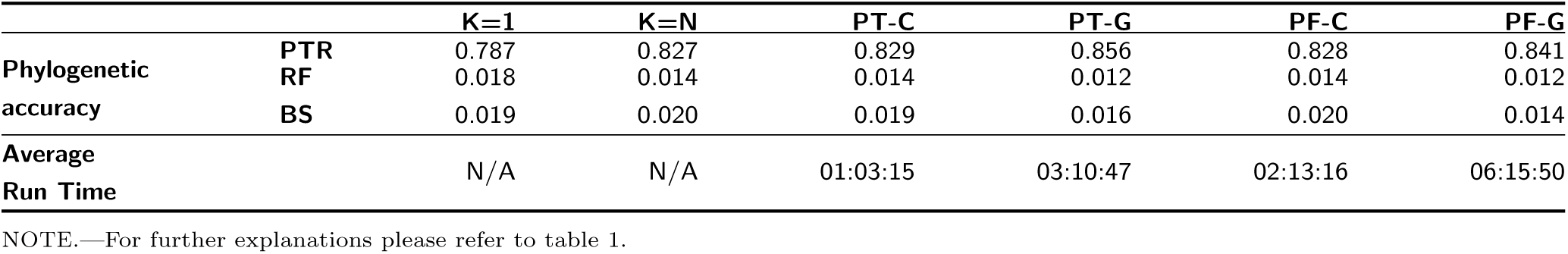
Phylogenetic accuracy for Simulation 2 (mosaic data blocks).

|  |  | K=1 | K=N | PT-C | PT-G | PF-C | PF-G |
| --- | --- | --- | --- | --- | --- | --- | --- |
| Phylogenetic accuracy | PTR | 0.787 | 0.827 | 0.829 | 0.856 | 0.828 | 0.841 |
|  | RF | 0.018 | 0.014 | 0.014 | 0.012 | 0.014 | 0.012 |
|  | BS | 0.019 | 0.020 | 0.019 | 0.016 | 0.020 | 0.014 |
| Average Run Time |  | N/A | N/A | 01:03:15 | 03:10:47 | 02:13:16 | 06:15:50 |

NOTE.—For further explanations please refer to table 1.

### Simulation 3: large-scale phylogenomic study

For large data sets (500,000-1,500,00 bp) the greedy algorithms can take very long, and only the hierarchical clustering algorithms were evaluated. In fact, even in this case PartitionFinder was not able to evaluate 20 out of the 200 replicates due to execution errors, while only 1 replicate failed for PartitionTest. All the comparisons in table 3 refer to the 180 replicates in common. The clustering algorithm implemented in PartitionTest (*PPR* = 0.056; *RI* = 0.989) was more accurate than its analog in PartitionFinder (*PPR* = 0.011; *RI* = 0.846) finding the true partitioning scheme, and 6.2 times faster on average (table 3). Both algorithms overestimated the true number of partitions, specially in the case of PF-C. The K=T and K=N *a priori* strategies always recovered the true topology, while K=1 failed in a few occasions. The average number of distinct topologies per replicate across methods was 1.05. Using the true partitioning scheme (K=T) resulted in very accurate branch lengths. The phylogenetic accuracy of PartitionTest and PartitionFinder was also very high, with the former providing slightly better branch length estimates and the latter finding the true topology in one additional replicate. Overall, PartitionTest was *∼*7 times faster than PartitionFinder.

**Table 3.**
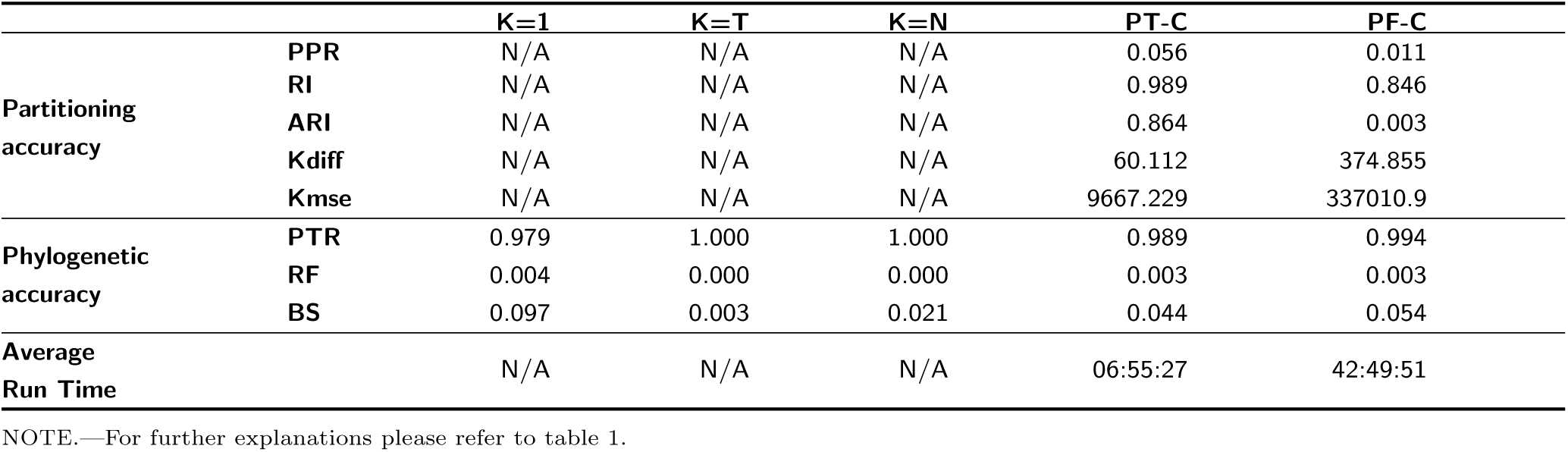
Partitioning and phylogenetic accuracy for Simulation 3 (phylogenomics).

### Simulation 4: rate variation

In the presence of rate variation among lineages and partitions the true partitioning scheme was never found (*PPR* = 0), although the RI scores were still high (*RI* = 0.932−0.953) (table 4). PartitionTest was slightly more accurate than PartitionFinder (*∼* 0.95 vs *∼* 0.93), and the accuracy of the best-fit partitioning schemes did not change when assuming independent vs. proportional branch lengths. The ARI values for PartitionFinder were low or very low. Both PartitionTest and PartitionFinder overestimated the true number of partitions, specially in the latter case. The different strategies showed very similar phylogenetic accuracy, with K=1 doing slightly worse. The true topology was found in most occasions (*P T R* = 0.895−0.907 for most strategies except for K=1, with *P T R* = 0.839), with small RF distances (0.005 for all strategies except PT-C-p and K=1, with RF=0.009) and similar BS (*∼* 0.428 for all strategies except PT-C- p and K=1, with BS=0.483). The average number of distinct topologies per replicate across methods was 1.25. Note that the branch length estimates were much worse than in previous scenarios in which the simulated branch lengths were the same across partitions. As expected, assuming proportional branch lengths resulted in faster run times. Overall, PartitionTest was *∼* 4 times faster than PartitionFinder.

**Table 4.** Partitioning and phylogenetic accuracy for Simulation 4 (rate variation).

|  |  | K=1 | K=T | K=N | PT-C | PT-G | PT-C-p | PT-G-p | PF-C | PF-G | PF-C-p | PF-G-p |
| --- | --- | --- | --- | --- | --- | --- | --- | --- | --- | --- | --- | --- |
| Partitioning accuracy | PPR | N/A | N/A | N/A | 0 | 0 | 0 | 0 | 0 | 0 | 0 | 0 |
|  | RI | N/A | N/A | N/A | 0.952 | 0.91 | 0.953 | 0.906 | 0.932 | 0.908 | 0.935 | 0.9 |
|  | ARI | N/A | N/A | N/A | 0.539 | 0.529 | 0.441 | 0.506 | 0.007 | 0.527 | 0.005 | 0.455 |
|  | Kdiff | N/A | N/A | N/A | 7.239 | -15.564 | 9.677 | -12.141 | 17.331 | -17.638 | 18.826 | -11.308 |
|  | Kmse | N/A | N/A | N/A | 73.269 | 307.855 | 120.923 | 201.002 | 378.886 | 386.877 | 428.989 | 179.385 |
| Phylogenetic accuracy | PTR | 0.839 | 0.907 | 0.904 | 0.895 | 0.866 | 0.903 | 0.888 | 0.904 | 0.865 | 0.904 | 0.888 |
|  | RF | 0.009 | 0.005 | 0.005 | 0.005 | 0.007 | 0.009 | 0.005 | 0.005 | 0.007 | 0.005 | 0.005 |
|  | BS | 0.483 | 0.429 | 0.428 | 0.428 | 0.43 | 0.483 | 0.431 | 0.428 | 0.431 | 0.428 | 0.43 |
| Average Run Time |  | N/A | N/A | N/A | 00:19:49 | 07:16:42 | 00:15:15 | 01:22:34 | 01:26:07 | 10:38:39 | 01:01:55 | 06:07:45 |

NOTE.—PT-C-p and PF-C-p assume that branch lengths are proportional across partitions. For further explanations please refer to table 1.

### Simulation 5: the effect of the likelihood optimization threshold

Changing the ML optimization threshold did not have a noticeable impact on the final inferences but dramatically influenced the running times. Higher epsilon thresholds (i.e., less thorough optimization) did not seem to influence much the resulting optimal partitioning schemes (figure 1a-c) or the resulting trees (with identical inferred topologies and very similar branch length estimates). However, the partitioning search algorithm was up to 4 times faster on average in this case. (figure 1d).

**FIG. 1.**
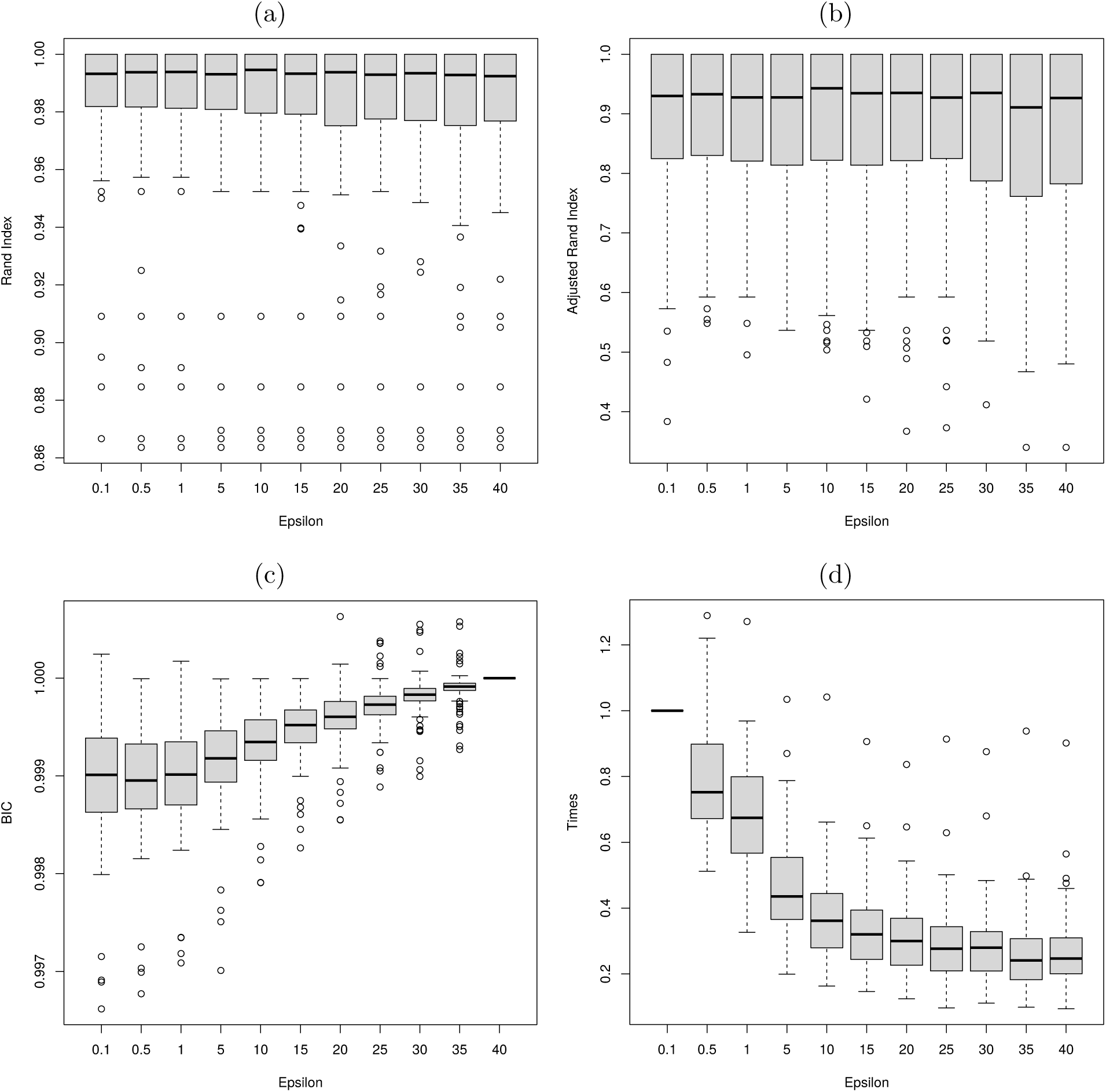
Partitioning sensitivity and running times as a function of the optimization threshold epsilon. (a) Rand Index. (b) Adjusted Rand Index. (c) Relative BIC scores (normalized to epsilon = 40) (d) Relative execution times (normalized to epsilon = 0.1).

### Analysis of real data

The optimal partitioning schemes identified in the real datasets were often different depending on the exact implementation (program and method) used, and in most cases without particularly obvious trends. With some exceptions, the assumption of proportional branch lengths across partitions resulted in more partitions in the optimal partitioning scheme (table S1). The number of model parameters in the optimal partitioning schemes was very variable. The greedy algorithms resulted in more or less partitions in the optimal partitioning scheme than the clustering algorithms depending on the data set. To make a legit comparison of the optimal BIC (Schwarz, 1978) scores found by the different algorithms we recomputed all the BIC scores in RAxML (Stamatakis, 2006) assuming proportional branch lengths (BIC*). No significant or consistent differences were observed. Regarding running times, the hierarchical clustering algorithms were clearly faster than the greedy algorithms, while assuming proportional branch lengths further reduced the computation time. On average, PartitionTest was times faster than PartitionFinder. The optimal partitioning schemes found by the different algorithms were often quite distinct (table S2), being most similar in general when the same algorithm but a different program was used (e.g., PartitionTest greedy vs. PartitionFinder greedy). The assumption of proportional/independent branch lengths across partitions often resulted in quite different partitioning schemes, with a slightly bigger influence than the program or algorithm used (table S3).

The ML trees estimated under the best-fit partitioning schemes found by the different methods were more or less distinct depending on the specific data set, with RF values ranging from 0 to 0.37. For data sets like *Endicott* or *Li*, the topological differences were highest, in particular regarding the assumption of proportional/independent branch lengths across partitions, which had the most noticeable effect across all data sets. For data sets like *Fong* all trees estimates were very similar, independently of the partitioning selection strategy. The branch scores were very low in practically every case, suggesting in principle that branch length estimates were not affected by the partitioning strategy, although we should note that in some cases the tree length was very small –like for *Endicott* – preventing large BS scores.

In addition, we also compared the ML trees found under the optimal partitioning schemes or under *a priori* partitioning scheme with a single partition (K=1), against the ML trees inferred using the K=N strategy (each partition assumed to evolve under an independent GTR+G model) (table 5). Again, the different strategies often resulted in different trees, except for the *Fong* data set, where all but one of the strategies resulted in the same topology.

**Table 5.** Comparison of the RAxML trees inferred under the optimal partitioning schemes found by different strategies in the 6 empirical data sets in the study.

| RF scores | Endicott | Fong | Hackett | Kaffenberg | Li | Wainwright |
| --- | --- | --- | --- | --- | --- | --- |
| **K=1** | 0.29 [tree A] | 0.00 [tree A] | 0.08 [tree A] | 0.13 [tree A] | 0.17 [tree A] | 0.09 [tree A] |
| **PT-C** | 0.29 [tree C] | 0.00 [tree A] | 0.08 [tree A] | 0.19 [tree B] | 0.03 [tree C] | 0.09 [tree A] |
| **PT-G** | 0.29 [tree A] | 0.07 [tree B] | 0.08 [tree C] | 0.07 [tree D] | 0.03 [tree E] | 0.05 [tree D] |
| **PT-C-p** | 0.36 [tree B] | 0.00 [tree A] | 0.11 [tree B] | 0.19 [tree B] | 0.24 [tree B] | 0.12 [tree B] |
| **PT-G-p** | 0.05 [tree D] | 0.00 [tree A] | 0.11 [tree B] | 0.05 [tree C] | 0.00 [tree D] | 0.01 [tree C] |
| **PF-C** | 0.29 [tree C] | 0.00 [tree A] | 0.08 [tree A] | 0.19 [tree B] | 0.17 [tree A] | 0.09 [tree A] |
| **PF-G** | 0.29 [tree A] | 0.00 [tree A] | 0.04 [tree F] | 0.07 [tree D] | 0.07 [tree G] | 0.07 [tree G] |
| **PF-C-p** | 0.12 [tree E] | 0.00 [tree A] | 0.02 [tree D] | 0.19 [tree B] | 0.01 [tree F] | 0.03 [tree E] |
| **PF-G-p** | 0.05 [tree F] | 0.00 [tree A] | 0.02 [tree E] | 0.13 [tree A] | 0.01 [tree F] | 0.00 [tree F] |
| **Num. different topologies** | 6/9 | 2/9 | 6/9 | 4/9 | 7/9 | 7/9 |

NOTE.—Cells contain the normalized RF scores against the RAxML trees assuming and independent GTR+G model per data block (K=N). Letters in brackets indicate the different topologies found.

## Discussion

### Identifying optimal partitioning schemes

Identifying the optimal partitioning scheme is not an easy task, but the different selection strategies studied here seem to perform quite well. In our simulations the exact true partitioning scheme was recovered up to 30% of the time when the number of data blocks was relatively small, and up to 5% of the time when this number was larger, which is still a remarkable result given the vast number of potential solutions. In the presence of rate variation among lineages and partitions the problem becomes much harder and the true partitioning scheme was never found. Still, most methods were able to identify most of the time and in most conditions optimal partitioning schemes closely related to the true partitioning scheme, as judged by the generally high Rand Index (RI) scores. Now, we should consider that the RI works a little bit different than what we might intuitively think. This index considers pairs of data blocks that belong to the same partition in both partitioning schemes, but also those belong to different partitions in the two schemes being compared. In contrast, our perception might be that a similarity measure between two partitioning schemes should count only those data blocks belong to the same partition in both schemes. Thus, some implementations like the clustering algorithm of PartitionFinder, that significantly overestimates the number of partitions, can still display high RI scores because many pairs of data blocks will belong to different partitions in both partitioning schemes. However, in this case the Adjusted RI scores were very low, highlighting the problem. Also, note that the RI does not consider the particular substitution models assigned to each partition. Hence, as long as two data blocks are grouped together, they are assumed to belong to the same partition, even if the best-fit models assigned to their partitions are different. The performance of the greedy and clustering algorithms was slightly different. Greedy algorithms tended to underestimate the number of partitions while the hierarchical clustering algorithms usually overestimated it. On average, greedy algorithms identified the true partitioning scheme more frequently than the clustering ones, but selected slightly less accurate partitioning schemes. Nevertheless, the computational complexity of the hierarchical clustering algorithms is much lower than that of the greedy algorithms (linear versus quadratic), so they were several times faster. Even though one might think of the schemes evaluated by the clustering algorithms as strict subsets of those evaluated by the greedy algorithm, this is not necessary the case. In a particular iteration where the initial partitioning scheme is the same for both algorithm it is true that hierarchical clustering inspects a subset of the candidate partitioning schemes in the greedy algorithm. However, the selected partitioning in that iteration scheme might not be the same for both algorithms, and hence from there they can take different paths that might result in different probabilities of getting stuck in local maxima. Therefore, the greedy algorithms do not necessarily reach always a better partitioning scheme than the clustering algorithms.

### Effect of partitioning on phylogenetic accuracy

In the simulations the different partitioning methods resulted in the inference of the same ML tree topology, with the only exception of the single partition strategy, which led to a different topology in some cases. Phylogenetic accuracy was quite high overall, even under rate variation among lineages and/or among partitions, or when there was obligate model misspecification (i.e., with mosaic data blocks). Although the greedy and hierarchical clustering strategies did not return the true partitioning scheme in most occasions, they still resulted in practically the same trees as those obtained under the true partitioning scheme. Only when a single partition was assumed *a priori* (i.e., the data was left unpartitioned), phylogenetic accuracy dropped down to some extent, up to 10% when the number of data blocks was not very large. This is in concordance with a previous simulation study that suggested that underpartitioning could negatively affect phylogenetic estimates under a Bayesian framework (Brown and Lemmon, 2007). However, while in the Bayesian case overly complex partitioning schemes also had some effect on posterior probabilities, in our simulations the ML trees did not seem to be much sensitive to overparameterization, an observation already made with real data by Li *et al.* (2008). In general, it is well known that ML phylogenetic estimation can be very robust to model misspecification when the phylogenetic problem is not too complex (e.g., Sullivan and Joyce, 2005) and this might at least partially explain why in the simulations ML phylogenetic estimation seems quite robust also to the partitioning scheme chosen and subsequent model assignment, despite for example having introduced branch length variation in the simulated trees or used mosaic data blocks.

Remarkably, the *a priori* assigning of independent GTR+G model to each data block led to very similar trees than the greedy and hierarchical clustering strategies, which advocates this as a very convenient strategy for the analysis of phylogenomic datasets. An alternative *a priori* K=N option not evaluated here might have been the independent assignment of best-fit models, for example identified with jModelTest (Darriba *et al.*, 2012), to each data block. While this is an obvious strategy, it was not explored here mainly because it cannot be currently implemented in RAxML, difficulting then a fair comparison among approaches. In any case, this strategy would require much more computation than the assignment of independent GTR+G models to each data block, which already results in optimal performance –practically the same as when using the true partitioning scheme.

Phylogenetic accuracy was almost perfect when the number of data blocks was very large, even for the single partition case. This is the expected behaviour because in our simulations all the data blocks were evolved under the same tree, so there was no phylogenetic incongruence among the different partitions. In real life phenomena like incomplete lineage sorting, gene duplication and loss and horizontal gene transfer can lead to a different scenario in which different partitions evolve under different gene trees, embedded within a single species tree (e.g., Martins *et al.* 2014). Thus, further studies could focus on evaluating the impact of partitioning on the inference of species trees.

### Empirical data analysis

The analysis of real data can be very helpful to show the relative fit of the different partitioning schemes and/or the congruence of the different phylogenetic estimates derived from them, beyond the simplicity of simulated data. On the other hand, in this case neither true partitioning scheme nor the true phylogeny is known, and accuracy cannot be directly measured. However, in our analyses of the six empirical data sets we saw more topological variation than in the simulations. In this case the partitioning selection strategy had a stronger effect than in the simulations, and the final tree estimates varied more depending on the method chosen. Nevertheless, this was not true for every dataset, and in all cases there was at least one selection strategy that resulted in the same tree as the unpartitioned scenario. These results are in agreement with previous empirical studies in which different partitioning schemes sometimes resulted in different trees, but also where the main differences were also observed when the data was left unpartitioned (Brandley *et al.*, 2005; Kainer and Lanfear, 2015; Leavitt *et al.*, 2013; Powell *et al.*, 2013; Ward *et al.*, 2010). For example, Kainer and Lanfear analyzed 34 data sets with different partitioning strategies that half or more of the time resulted in different trees, albeit the differences among them were not significant, with average RF distances smaller than 10%, except for the unpartitioned case, which implied significant differences. The same was true for branch lengths and bootstrap values, were only the use of a single partition made a difference in some cases..

### Alternative partitioning approaches

The definition of homogeneous data blocks can be a problem under certain circumstances, like in the case of non-coding regions, or when there is significant heterogeneity at a local scale. However, in our (simple) simulation of non-homogeneous data blocks, phylogenetic accuracy was still reasonably high. Wu *et al.* (2013) described an elegant Bayesian framework in which the partitioning scheme is treated as a random variable using a Dirichlet process prior (DPP). This method is able to simultaneously identify the number of partitions, their best-fitting models, and assign sites to each one of them. While this approach is certainly much more flexible than previous strategies, explicitly considers the uncertainty associated with partitioning and improves model fit, it has not been demonstrated yet to lead to more accurate trees. Unfortunately, its heavy computational requirements, and the restriction of only sites being the units if the assignment seem to have limited for now its widespread application to real data. Indeed, it might be very interesting to see a DPP method-in fact a special case of the one just described- that works with user-defined data blocks. Very recently, Frandsen *et al.* (2015) introduced a promising algorithm for phylogenetic partitioning that uses rates of evolution instead of substitution patterns, also avoiding the need for an arbitrary delimitation of data blocks. While this method can increase model fit, again its advantage over data block methods like those studied here has not been demonstrated in terms of phylogenetic accuracy.

### PartitionTest vs. PartitionFinder

In the majority of the conditions explored in the simulations PartitionTest was slightly more accurate than PartitionFinder both regarding the identification of optimal partitioning schemes and tree estimation. Although these differences were small, they were consistent. Importantly, PartitionTest is much faster than PartitionFinder, between 1.5 and 7 times faster, in particular with large data sets. PartitionFinder is implemented in Python and delegates the phylogenetic calculations on external third party software, like PhyML (Guindon and Gascuel, 2003) or RAxML. On the other hand, PartitionTest is implemented in C++ and keeps a finer control over the phylogenetic calculations through the use use of the PLL (Flouri *et al.*, 2015).

## Conclusions

Several strategies for the selection of best-fit partitioning schemes have been recently introduced for the phylogenetic analysis of multilocus data sets. Here we evaluated different partitioning approaches using comprehensive computer simulations and real data. We conclude that hierarchical clustering algorithms should be preferred over existing greedy algorithms for the selection of best-fit partitioning scheme, because under the conditions explored, they were much faster with practically the same accuracy. However, our simulations also suggest that ML phylogenetic inference is quite robust to the partitioning scheme, at least as far as single models are assigned to the final partitions using a statistical procedure. In this case, any reasonable partitioning scheme seems to perform reasonably well, including the *a priori* assignment of GTR+G model to each data block. To be on the safe side, leaving the data unpartitioned should be avoided.

## Materials and methods

### Partitions and partitioning schemes

Let us consider set of aligned nucleotide or amino acid sequences of any length (the “data”). Following Lanfear *et al.* (2012), we define “data block” as a set of alignment sites that are assumed to evolve in a similar way, normally user-defined. Typical examples of data blocks in phylogenetics are amplified gene fragments, gene families, assembled loci from NGS reads, introns/exons, or sets of codon positions. A “partition” (“subset” in Lanfear *et al.* (2012) will be a set of one or more particular data blocks. A partition can be made of, for example, a single gene fragment, family or locus, multiple gene fragments, families or loci, or consist of the set of all 1st and 2nd codon positions in an alignment. Finally, a set of non-overlapping partitions that cover the whole alignment will be called a “partitioning scheme”. The partitioning problem consists of, given a predefined set of data blocks, finding the optimal partitioning scheme for a given alignment. In our case, we want to optimize the partitions with regard to the assignment of substitution models. Note that a “model” here will be a particular model of nucleotide substitution or amino acid replacement together with parameter values. That is, K80 (*ti/tv* = 2) would be a different model than K80+G or JC, but also than K80 (*ti/tv* = 8). For example, if we have a multilocus alignment with, say 100 concatenated genes, our aim is to find out whether we should use 100 different substitution models, just one (all genes evolving under exactly the same model), or something in between, in which case we would need to assign 2-99 models to the 100 genes. Note that there are two related questions here, which are: (i) how many different models should we use (i.e., the number of partitions), and (ii) which partitions evolve under which model (i.e., the partitioning scheme). In general, given *n* initial data blocks, the number of possible partitioning schemes, *B*(*n*), is given by the Bell numbers, which are the sum from 1 to *n* of the Stirling numbers of the second kind, *S*(*n,k*), where *k* is the number of partitions:

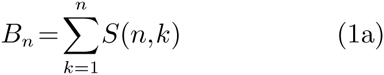

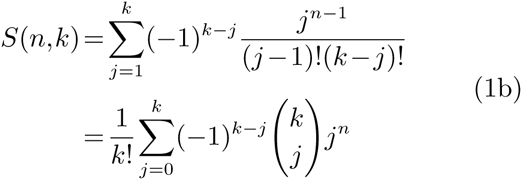

The number of partitioning schemes grows very quickly. For example, for 4 data blocks there are 15 different partitioning schemes, but for 20 there are already 5.8×10^12^. Clearly, finding the optimal partitioning scheme and assigned models is a very intensive task, and rapidly becomes computationally unfeasible.

### Selecting optimal partitioning schemes

In order to select optimal partitioning schemes at the phylogenomic level, we have implemented *de novo* (in the program PartitionTest, see below) a set of heuristic algorithms that are very similar to those already available in the software PartitionFinder. The main steps in these algorithms are:

1. Estimate an initial tree.
2. Define a set of candidate partitioning schemes.
3. Select the best-fit substitution/replacement model for each partition.
4. Compute the score of each partitioning scheme and select the best one accordingly.
5. Return to step 2 until there is no score improvement or until the current partitioning scheme includes a single partition.

#### Step 1. Initial tree estimate

The starting tree topology can be user-defined or estimated using a particular phylogenetic method.

#### Step 2. Define a set of candidate partitioning schemes

The initial partitioning scheme is the set of data blocks defined by the user. For each iteration, new partitioning schemes are proposed as potentially better candidates, given the currently best partitioning scheme, and using a greedy or a hierarchical clustering algorithm.

The greedy algorithm defines 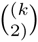 candidate partitioning schemes of size (*k* – 1) by merging all possible pairs of partitions, where k is the number of partitions in the current best partitioning scheme. This algorithm is identical to the greedy algorithm implemented in PartitionFinder. Its computational complexity is *O*(*n*^2^), so if the number of initial partitions (*n*) is large the required computational time will be considerable.

The hierarchical clustering algorithm defines *r* candidate partitioning schemes of size (*k* – 1) by merging the *r* closest pairs of partitions, given a matrix of pairwise distances between partitions (*D*(*m*_*i*_,*m*_*j*_), see below). The parameter *r* is defined by the user. A strict hierarchical clustering (i.e., *r* = 1) will evaluate a maximum of n candidate partitioning schemes. Although *r* can be defined in the range (0,*∞*), only a maximum of *n*(*n-*1)*/*2 new candidate partitioning schemes can be proposed, so if *r ≥ n*(*n* – 1)*/*2, this algorithm will behave exactly as the greedy algorithm. The pairwise distances between partitions *i* and *j* are calculated using the maximum likelihood estiamtes (MLEs) of the best-fit substitution model parameters for each partition:

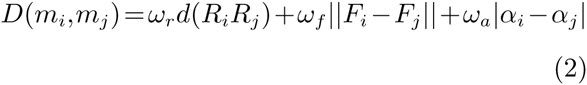

where *R* = *{r_ac_,r_ag_,r_at_,r_cg_,r_ct_,r_gt_}* are the substitution rates, *F* = *{f_a_,f_c_,f_g_,f_t_}* are the base frequencies, and *α* is the alpha shape for the gamma rate variation among sites (+G). Because the substitution rates are usually estimated relative to each other, we first scale them such that their euclidean distance is minimized:

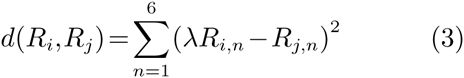

Deriving this function we obtain:

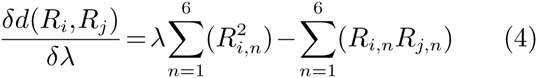

whose minimum is located at:

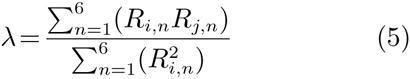

We include different weights (*ω*_*r*_, *ω*_*f*_, *ω*_*a*_ and *ω*_*p*_) for each part of the distance formula, that the user can specify. By default these values are set to those that maximized accuracy (finding the true partitioning scheme) in pilot simulations. Note that the hierarchical clustering algorithm implemented PartitionFinder specifies a slightly different formulae than PartitionTest for the distance calculation. The computational complexity of the hierarchical clustering algorithm is *O*(*rn*), so the required computational time should be affordable even for very large data sets (e.g., with *>* 1,000 initial partitions).

#### Step 3. Select a substitution model for each partition

For each partition, likelihood scores are calculated given a tree and a model of substitution/replacement. Best-fit substitution/replacement models with associated parameter values are then identified using the Akaike Information Criterion (AIC, Akaike, 1973), corrected AIC (AICc, Sugiura, 1978), Bayesian Information Criterion (BIC, Schwarz, 1978) or Decision Theory (DT, Minin *et al.*, 2003). Alternatively, a fixed substitution/replacement model can be assigned for every partition, with unlinked parameter values that are independently optimized. For the likelihood calculations the tree topology can be fixed (i.e., the starting tree topology is used for every calculation) or reoptimized using maximum likelihood for each partition. Branch lengths across partitions can be assumed to be independent for each partition (unlinked) or proportional among partitions (linked). In the independent model the branch lengths are reoptimized for every new partition. In the proportional model a set of global branch lengths is estimated at the beginning for the whole data set, with a scaling parameter being optimized for every new partition.

#### Step 4. Compute the score of each partitioning scheme

The score of a partitioning scheme will be calculated in two different ways depending on the occurrence of linked/unlinked branch lengths.

If branch lengths are unlinked across partitions, model parameters and branch lengths are optimized independently for each partition.

Therefore, the BIC score of a partitioning scheme is simply the sum of the individual scores of its partitions:

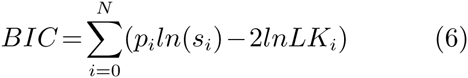

where *p*_*i*_ is the number of parameters, *s*_*i*_ is the sample size, and *LK*_*i*_ is the likelihood score of partition *i*.

However, if branch lengths are linked proportionally across partitions, the score of the partitioning scheme with linked parameters is computed as follows:

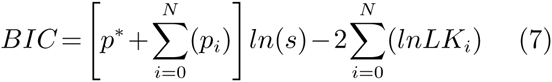

where *p*_*i*_ is the number of parameters of partition *i*, *p*^*\**^ is the number of parameters globally optimized for the partitioning scheme, and *s* is the sample size of the entire partitioning scheme.

### PartitionTest software

We have implemented the algorithms described above in the program PartitionTest, available from *https://github.com/ddarriba/partitiontest* The greedy search algorithm in PartitionTest is essentially the same as the one implemented in PartitionFinder, but the hierarchical clustering algorithm uses slightly different distances.

PartitionTest makes an intensive use of the Phylogenetic Likelihood Library (PLL) for carrying out all likelihood computations, including tree estimation. The PLL speeds up the calculations considerably. During ML estimation of model parameters and trees with PLL, a parameter *E* regulates how thorough are the mathematical optimizations. Basically, epsilon is a numerical threshold under which the improvement in likelihood is considered not worthy and the optimization stops. When epsilon is small the optimization is more thorough and takes more time. PartitionTest tests implements several computational strategies to avoid repeated calculations, including checkpoint and restarting capabilities, allowing its use in systems with per-job time restriction, like many High Performance Computing (HPC) clusters. In order to choose the best substitution/replacement model for each partition, PartitionTest considers 22 models of DNA substitution (the +G models in jModelTest2) and 36 empirical models of amino acid replacement (the same as RAxML excluding LG4M and LG4X). All of them assume rate heterogeneity among sites using a discrete gamma distribution with four categories. If desired, PartitionTest is also able to estimate ML trees from the optimal partitioning scheme.

### Benchmarking of partitioning algorithms

We devised a set of experiments with simulated and real DNA sequence data to compare different partitioning strategies. The main questions asked were (i) how accurate (close to truth) are the optimal partitions identified by the different algorithm, and (ii) what is the impact of the different partitioning strategies on phylogenetic accuracy. The different partitioning strategies evaluated were three *a priori* partitioning schemes plus different algorithms implemented in PartitionTest and PartitionFinder:

1. A single partition, or unpartitioned (K=1)
2. One partition for each data block (K=N)
3. The simulated partitioning scheme (K=T)
4. PartitionTest hierarchical clustering with independent branch lengths across partitions (PT-C)
5. PartitionTest greedy with independent branch lengths across partitions (PT-G)
6. PartitionTest hierarchical clustering with proportional branch lengths across partitions (PT-C-p)
7. PartitionTest greedy with proportional branch lengths across partitions (PT-G-p)
8. PartitionFinder greedy with independent branch lengths across partitions (PF-G)
9. PartitionFinder hierarchical clustering with independent branch lengths across partitions (PF-C)
10. PartitionFinder hierarchical clustering with proportional branch lengths across partitions (PF-C-p)
11. PartitionFinder greedy with proportional branch lengths across partitions (PF-G-p)

Strategies 6-7 and 10-11 were only evaluated in Simulation 4 (see below). All the analyses were carried out in a computer with 2 hexa-core Intel Xeon X5675 @ 3.07GHz processors (12 cores) and 50GB of memory, with Hyper-threading disabled. We used a single core per run to facilitate running time comparisons.

#### Computer simulations

The first four experiments consisted of a series of computer simulations aiming to recreate different biological scenarios, while the last one was designed to assess the sensitivity of the results to the level of parameter optimization (table 6). In our simulations, parameter values were not fixed along a grid but sampled from predefined statistical distributions, allowing us to explore a large parameter space and to carry out *ad hoc* analyses of the results.

- Simulation 1: with a limited number of data blocks, typical of a multi-gene phylogenetic study.
- Simulation 2: with pairs of data blocks merged at random before the analysis. Our intention was to represent an scenario where sites inside data blocks did not evolve in an homogeneous fashion, as assumed by definition. Instead, in this simulation data blocks are mosaics of two distinct evolutionary processes.
- Simulation 3: with a large number of data blocks, typical of a large-scale phylogenomic study.
- Simulation 4: with rate variation among partitions and lineages. In this case the branch lengths for each partition were scaled using two random multipliers. A global multiplier *∼*U(0.25, 4) was applied to all branches, while a local multiplier *∼*U(0.8, 1.2) was chosen for each branch. For the analysis of the simulated data, we used both the independendent and proportional branch length models.

**Table 6.** Simulation summary. Parameter values were chosen to reflect a range of plausible biological scenarios.

|  | Sim1 | Sim2 | Sim3 | Sim4 | Sim5 |
| --- | --- | --- | --- | --- | --- |
| **N, number of genes** | U(10,50) | U(5,25) | 1000 | U(10,50) | U(5,50) |
| **K, number of partitions** | U(1,N) | NA | U(1,N) | U(1,N) | U(1,N) |
| **Gene length** | U(500,1500) | U(1000,3000) | U(500,1500) | U(500,1500) | U(500,1500) |
| **Number of taxa** | U(6,40) | U(6,40) | U(6,40) | U(6,40) | U(8,140) |
| **Topology** | Fixed | Fixed | Fixed | Branch Length multiplier | Fixed |
| **Number of replicates** | 4,000 | 4,000 | 200 | 1,000 | 100 |
| **Tree length** | U(0.5,15) | U(0.5,12) | U(0.5,12) | U(0.5,12) | U(0.5,12) |

Simulation 5: here we tested the impact of the optimization threshold epsilon on the resulting partitioning schemes and topologies, in order to find a good compromise between computational time and accuracy.

For each replicate the simulation proceeded as follows:

1. *N* data blocks were generated according to U[10,50] with variable lengths chosen from U[500,1500].
2. Data blocks were randomly assigned to *K* partitions, where *K ∼*U[1,N].
3. Each partition was assigned a random model of nucleotide substitution.
  a. A model family (M) is chosen from the 22 nucleotide substitution model families *∼*U(0,21).
  b. A model of rate variation was chosen among 4 possibilities *∼*U(0,3): no rate variation (M), including a proportion of invariable sites (M+I), including gamma rate variation among sites (+G), and including both a proportion of invariable sites and gamma rate variation among sites (+I+G).
4. Specific model parameter values were chosen from prior distributions.
  a. Nucleotide frequencies: equal or *∼*Dirichlet(1.0,1.0,1.0,1.0).
  b. Transition/transversion rate: *∼*Gamma(2,1) truncated between 2 and 10
  c. R-matrix parameters *∼*Dirichlet(6,16,2,8,20,4) scaled with the last rate (taking free parameters as necessary).
  d. Proportion of invariable sites *∼*Beta(1,3) truncated between 0.2 and 0.8.
  e. Gamma shape for rate variation among sites *∼*Exponential(2) truncated between 0.5 and 5.
5. A random non-ultrametric rooted tree topology with number of taxa *∼*U(6,40) and branch lengths *∼*Exponential(1,10) was simulated with the function rtree from the ape package (Paradis *et al.*, 2004) in R. The total tree length is scaled so tree length *∼*U[2, 12].
6. Each partition was evolved under this tree according to the chosen substitution model parameters using INDELible (Fletcher and Yang, 2009), resulting in a multiple sequence alignment.
7. Optimal partitions were identified from this alignment according to the partitioning strategies listed above, using the default settings in each software. See Lanfear *et al.* (2012) for details.
8. A ML tree was estimated from this alignment according to the optimal partitioning schemes identified under each strategy, using RAxML.

#### Analysis of real data

We also reanalyzed some of the real datasets previously used in the evaluation of PartitionFinder (table 7). As in the simulations, optimal partitioning schemes were selected under the different partitioning strategies evaluated, and used to infer ML trees with RAxML.

**Table 7.** Description of the empirical datasets evaluated in this study.

| Short name | Clade | Number of taxa | Sequence length | Number of data blocks | Average number of sites per data block | Reference |
| --- | --- | --- | --- | --- | --- | --- |
| Endicott | Humans (Homo sapiens) | 179 | 13857 | 42 | 329.92 | Endicott and Ho (2008) |
| Fong | Vertebrates (Vertebrata) | 16 | 25919 | 168 | 154.28 | Fong et al. (2012) |
| Hackett | Birds (Aves) | 171 | 52383 | 168 | 277.16 | Hackett et al. (2008) |
| Kaffenberger | Frogs (Gephyromantis) | 54 | 6145 | 27 | 277.59 | Kaffenberger et al. (2012) |
| Li | Ray-finned fishes (Actubioterygii) | 56 | 7995 | 30 | 266.5 | Li et al. (2008) |
| Wainwright | Ray-finned fishes (Acanthomorpha) | 188 | 8439 | 30 | 281.3 | Wainwright et al. (2012) |

#### Evaluation of partitioning and phylogenetic accuracy

*Partitioning accuracy* In order to compare the selected partitioning schemes obtained under the different partitioning strategies with the true partitions (simulation), or among themselves (real data), we computed different statistics. We counted how many times the exact true partitioning scheme was identified (PPR = Perfect Partitioning Recovery). We also calculated the Rand Index (Rand, 1971) (RI), a measure of the similarity between two clusterings that is constructed as follows. Given a set of *n* data blocks *S* = *{o*_1_,*…,o_n_}* and two partitioning schemes of *S* named *X* and *Y* with *r* and *s* partitions, respectively, *X* = *{X*_1_,*…,X_r_}* and *Y* = *{Y*_1_,*…,Y_s_}*, define the following:

- *a*, the number of pairs of data blocks in S that are in the same partition in *X* and in the same partition in *Y*.
- *b*, the number of pairs of data blocks in S that are in different partition in *X* and in different partition in *Y*.
- *c*, the number of pairs of data blocks in S that are in the same partition in *X* and in different partition in *Y*.
- *d*, the number of pairs of data blocks in S that are in different partition in *X* and in the same partition in *Y*.

Intuitively, *a*+*b* can be considered as the number of agreements between *X* and *Y* and *c*+*d* as the number of disagreements between *X* and *Y*. With these counts in place, the RI is computed as follows:

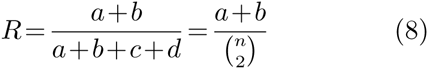

The RI is a value between 0 and 1, with 0 indicating that the two partitioning schemes are completely different and 1 that they are identical. In addition, we calculated the adjusted Rand index (Hubert and Arabie, 1985) (ARI), which measures the probability that a given RI was achieved by chance. The ARI can yield negative values if the observed RI is smaller than the expected RI. In this case the overlap between two partitioning schemes *X* and *Y* can be summarized in a contingency table where each entry *n*_*ij*_ denotes the number of data blocks in common between partition *X*_*i*_ and *Y*_*j*_, like this:

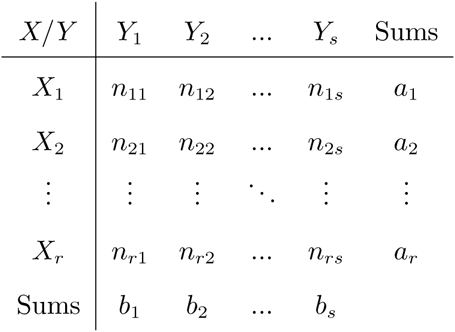

Then the ARI is calculated as:

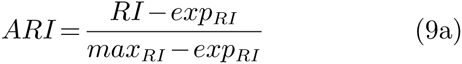

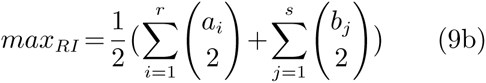

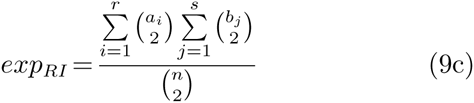

where *a*_*i*_ and *b*_*j*_ are values from the contingency table.

We also computed two statistics that reflect if the number of partitions is under or overestimated (Kdiff = average number of true partitions – number of partitions in the optimal partitioning scheme), and the mean square error of this deviation (Kmse).

#### Phylogenetic accuracy

In order to compare the inferred ML trees obtained under the different partitioning strategies with the true, generating trees (in the case of computer simulations), or among themselves (in the case of real data), we calculated: (i) how many times the exact true tree topology was identified (PTR = Perfect Topology Recovery), (ii) the Robinson-Foulds metric (RF) (Robinson and Foulds, 1981), that only considers the topology, and (iii) the branch score difference (BS) (Kuhner and Felsenstein, 1994), which takes also into account the branch lengths. In order to compare measurements from trees with different sizes, we scaled both the RF and BS so they were expressed per branch. We consider as outliers those simulation replicates that resulted in any BS difference (per-branch) higher than three. Even if the tree topologies were completely different, such a large BS distance could only be caused by a extremely long average branch length in one of the trees, suggesting an optimization error. This threshold resulted in only less than 1% of the replicates being treated as outliers.

## Supplementary Material

Supplementary tables S1–S3 are available at Molecular Biology and Evolution online (http://www.mbe.oxfordjournals.org/).

## Acknowledgments

This work was supported by the European Research Council (ERC-2007Stg 203161- PHYGENOM to D.P.) and the Spanish Government (research grant BFU2012-33038 to D.P.).

---

**Table S1.** Optimal partitioning schemes for the empirical datasets.

| **Endicott** | **PT-C** | **PT-G** | **PT-C-p** | **PT-G-p** | **PF-C** | **PF-G** | **PF-C-p** | **PF-G-p** |
| --- | --- | --- | --- | --- | --- | --- | --- | --- |
| **Npar** | 364 | 364 | 427 | 400 | 364 | 364 | 400 | 409 |
| **K** | 1 | 1 | 8 | 5 | 1 | 1 | 5 | 6 |
| **BIC\*** | 71409 | 71409 | 69439 | 69328 | 71409 | 71409 | 70193 | 69316 |
| **Run Time** | 00:02:22 | 00:26:30 | 00:01:09 | 00:16:53 | 00:04:17 | 00:55:42 | 00:03:41 | 00:46:44 |
| **Fong** |  |  |  |  |  |  |  |  |
| **Npar** | 497 | 128 | 182 | 164 | 128 | 110 | 218 | 137 |
| **K** | 52 | 11 | 17 | 15 | 11 | 9 | 21 | 12 |
| **BIC\*** | 285842 | 283542 | 284787 | 283508 | 293635 | 284146 | 285117 | 283358 |
| **Run Time** | 00:03:04 | 02:02:24 | 00:03:16 | 01:25:42 | 00:07:03 | 06:18:12 | 00:03:51 | 03:18:30 |
| **Hackett** |  |  |  |  |  |  |  |  |
| **Npar** | 348 | 366 | 1311 | 627 | 348 | 366 | 753 | 555 |
| **K** | 1 | 3 | 108 | 32 | 1 | 3 | 46 | 24 |
| **BIC\*** | 1862900 | 1844431 | 1841869 | 1836829 | 1862900 | 1844255 | 1843320 | 1837617 |
| **Run Time** | 04:11:04 | 72:52:40 | 00:41:17 | 13:17:33 | 04:23:35 | 81:28:25 | 01:35:34 | 52:32:22 |
| **Kaffenberger** |  |  |  |  |  |  |  |  |
| **Npar** | 132 | 141 | 186 | 204 | 114 | 132 | 168 | 186 |
| **K** | 3 | 4 | 9 | 11 | 1 | 3 | 7 | 9 |
| **BIC\*** | 130682 | 129538 | 129635 | 128791 | 128950 | 130169 | 131406 | 128955 |
| **Run Time** | 00:01:48 | 00:12:34 | 00:01:20 | 00:05:35 | 00:05:06 | 00:32:28 | 00:03:05 | 00:21:43 |
| **Li** |  |  |  |  |  |  |  |  |
| **Npar** | 190 | 145 | 280 | 208 | 163 | 127 | 217 | 199 |
| **K** | 9 | 4 | 19 | 11 | 6 | 2 | 12 | 10 |
| **BIC\*** | 257289 | 256878 | 256095 | 255849 | 260116 | 257749 | 256437 | 255787 |
| **Run Time** | 00:02:10 | 00:20:28 | 00:02:03 | 00:07:46 | 00:05:59 | 00:50:18 | 00:03:40 | 00:31:54 |
| **Wainwright** |  |  |  |  |  |  |  |  |
| **Npar** | 382 | 400 | 553 | 481 | 382 | 391 | 571 | 490 |
| **K** | 1 | 3 | 20 | 12 | 1 | 2 | 22 | 13 |
| **BIC\*** | 486930 | 480516 | 477486 | 477661 | 486930 | 480892 | 477932 | 477702 |
| **Run Time** | 00:09:14 | 01:23:36 | 00:09:05 | 00:29:15 | 00:25:19 | 03:25:53 | 00:13:49 | 01:59:03 |

NOTE.—For each dataset the table includes the total number of parameters in the optimal partitioning schemes (Npar), the number of partitions in the optimal partitioning schemes (K), the BIC scores of the optimal partitioning schemes assuming proportional branch lengths and recomputed in RaxML (BIC\*), and the time to select an optimal partitioning schemes (Run Time) in hours, minutes and seconds (hh:mm:ss).

**Table S2.** Rand-Index between the optimal partitioning schemes found under different strategies in the real datasets studied.

| Endicot |  |  |  |  |  |  |  |
| --- | --- | --- | --- | --- | --- | --- | --- |
|  | PT-C | PT-G | PT-C-p | PT-G-p | PF-C | PF-G | PF-Cp |
| PT-G | 1.00(0) |  |  |  |  |  |  |
| PT-C-p | 0.27(6) | 0.27(6) |  |  |  |  |  |
| PT-G-p | 0.21(5) | 0.21(5) | 0.78(-1) |  |  |  |  |
| PF-C | 1.00(0) | 1.00(0) | 0.27(-6) | 0.21(-5) |  |  |  |
| PF-G | 1.00(0) | 1.00(0) | 0.27(-6) | 0.21(-5) | 1.00(0) |  |  |
| PF-Cp | 0.29(4) | 0.29(4) | 0.88(-2) | 0.79(-1) | 0.29(4) | 0.29(4) |  |
| PF-G-p | 0.23(5) | 0.23(5) | 0.77(-1) | 0.97(0) | 0.23(5) | 0.23(5) | 0.77(1) |
| Fong |  |  |  |  |  |  |  |
|  | PT-C | PT-G | PT-C-p | PT-G-p | PF-C | PF-G | PF-C-p |
| PT-G | 0.79(-8) |  |  |  |  |  |  |
| PT-C-p | 0.8(4) | 0.83(4) |  |  |  |  |  |
| PT-G-p | 0.78(-6) | 0.93(2) | 0.8(-2) |  |  |  |  |
| PF-C | 0.67(6) | 0.73(14) | 0.71(10) | 0.73(12) |  |  |  |
| PF-G | 0.78(-11) | 0.90(-3) | 0.83(-7) | 0.87(-5) | 0.73(-17) |  |  |
| PF-C-p | 0.72(1) | 0.85(9) | 0.78(5) | 0.84(7) | 0.74(-5) | 0.84(12) |  |
| PF-G-p | 0.77(-7) | 0.91(1) | 0.82(-3) | 0.90(-1) | 0.74(-13) | 0.91(4) | 0.87(-8) |
| Hackett |  |  |  |  |  |  |  |
|  | PT-C | PT-G | PT-C-p | PT-G-p | PF-C | PF-G | PF-C-p |
| PT-G | 0.36(3) |  |  |  |  |  |  |
| PT-C-p | 0.06(55) | 0.68(52) |  |  |  |  |  |
| PT-G-p | 0.06(27) | 0.69(24) | 0.91(-28) |  |  |  |  |
| PF-C | 1.00(0) | 0.36(-3) | 0.06(-55) | 0.06(-27) |  |  |  |
| PF-G | 0.37(2) | 0.57(-1) | 0.62(-53) | 0.62(-25) | 0.37(2) |  |  |
| PF-C-p | 0.05(44) | 0.63(41) | 0.90(-11) | 0.91(17) | 0.05(44) | 0.65(42) |  |
| PF-G-p | 0.05(23) | 0.62(20) | 0.90(-32) | 0.90(-4) | 0.05(23) | 0.67(21) | 0.92(-21) |
| Kaffenberg |  |  |  |  |  |  |  |
|  | PT-C | PT-G | PT-C-p | PT-G-p | PF-C | PF-G | PF-Cp |
| PT-G | 0.47(1) |  |  |  |  |  |  |
| PT-C-p | 0.35(9) | 0.86(8) |  |  |  |  |  |
| PT-G-p | 0.38(7) | 0.90(6) | 0.96(-2) |  |  |  |  |
| PF-C | 0.79(-1) | 0.45(-2) | 0.34(-10) | 0.37(-8) |  |  |  |
| PF-G | 0.64(0) | 0.80(-1) | 0.67(-9) | 0.71(-7) | 0.57(1) |  |  |
| PF-C-p | 0.44(4) | 0.72(3) | 0.71(-5) | 0.73(-3) | 0.37(5) | 0.77(4) |  |
| PF-G-p | 0.49(5) | 0.70(4) | 0.71(-4) | 0.72(-2) | 0.36(6) | 0.73(5) | 0.73(1) |
| Li |  |  |  |  |  |  |  |
|  | PT-C | PT-G | PT-C-p | PT-G-p | PF-C | PF-G | PF-C-p |
| PT-G | 0.71(-11) |  |  |  |  |  |  |
| PT-C-p | 0.82(5) | 0.75(16) |  |  |  |  |  |
| PT-G-p | 0.74(-4) | 0.75(7) | 0.90(-9) |  |  |  |  |
| PF-C | 0.21(-14) | 0.28(-3) | 0.04(-19) | 0.09(-10) |  |  |  |
| PF-G | 0.67(-13) | 0.74(-2) | 0.50(-18) | 0.55(-9) | 0.54(1) |  |  |
| PF-C-p | 0.76(-3) | 0.72(8) | 0.86(-8) | 0.83(1) | 0.11(11) | 0.57(10) |  |
| PF-G-p | 0.76(-5) | 0.75(6) | 0.90(-10) | 0.86(-1) | 0.08(9) | 0.54(8) | 0.85(-2) |
| Wainwright |  |  |  |  |  |  |  |
|  | PT-C | PT-G | PT-C-p | PT-G-p | PF-C | PF-G | PF-C-p |
| PT-G | 0.50(2) |  |  |  |  |  |  |
| PT-C-p | 0.04(18) | 0.54(16) |  |  |  |  |  |
| PT-G-p | 0.06(11) | 0.56(9) | 0.94(-7) |  |  |  |  |
| PF-C | 1.00(0) | 0.50(-2) | 0.04(-18) | 0.06(-11) |  |  |  |
| PF-G | 0.54(1) | 0.96(-1) | 0.50(-17) | 0.52(-10) | 0.54(1) |  |  |
| PF-C-p | 0.03(21) | 0.52(19) | 0.94(3) | 0.94(10) | 0.03(21) | 0.49(20) |  |
| PF-G-p | 0.06(12) | 0.55(10) | 0.94(-6) | 0.94(1) | 0.06(12) | 0.52(11) | 0.93(-9) |

NOTE.—In brackets, the difference in the number of partitions (row - column).

**Table S3.** Average Rand-Index across data sets between PartitionTest and PartitionFinder, between hierarchical clustering and Greedy algorithms, and proportional versus independent branch lengths.

| **Dataset** | **PartitionTest**<br>**vs.**<br>**PartitionFinder** | **Clustering**<br>**vs.**<br>**Greedy** | **Proportional**<br>**vs.**<br>**Independent** |
| --- | --- | --- | --- |
| Endicott | 0.4693 | 0.5707 | 0.2515 |
| Fong | 0.8030 | 0.7941 | 0.8090 |
| Hackett | 0.4523 | 0.4935 | 0.3504 |
| Kaffenberger | 0.5858 | 0.6160 | 0.5716 |
| Li | 0.4514 | 0.6193 | 0.5346 |
| Wainwright | 0.4260 | 0.5074 | 0.2856 |
| **Average** | **0.5313** | **0.6002** | **0.4671** |

